# HAND2 is a novel obesity-linked adipogenic transcription factor regulated by glucocorticoid signaling

**DOI:** 10.1101/2020.05.18.102699

**Authors:** Maude Giroud, Foivos-Filippos Tsokanos, Giorgio Caratti, Sajjad Khani, Elena Sophie Vogl, Martin Imler, Christina Glantschnig, Manuel Gil-Lozano, Stefan Kotschi, Daniela Hass, Asrar Ali Khan, Marcos Rios Garcia, Frits Mattijssen, Adriano Maida, Daniel Tews, Pamela Fischer-Posovszky, Annette Feuchtinger, Kirsi A. Virtanen, Johannes Beckers, Martin Wabitsch, Matthias Blüher, Jan Tuckerman, Marcel Scheideler, Alexander Bartelt, Stephan Herzig

## Abstract

Adipocytes are critical cornerstones of energy metabolism. While obesity-induced adipocyte dysfunction is associated with insulin resistance and systemic metabolic disturbances, adipogenesis, the formation of new adipocytes and healthy adipose tissue expansion are associated with metabolic benefits. Understanding the molecular mechanisms governing adipogenesis is of great clinical potential to efficiently restore metabolic health in obesity. Here we show that Heart- and neural crest derivatives-expressed protein 2 (HAND2) is an obesity-linked adipocyte transcription factor regulated by glucocorticoids and required for adipocyte differentiation *in vitro*. In a large cohort of humans with obesity, white adipose tissue (WAT) *HAND2* expression was correlated to body-mass-index (BMI). The *HAND2* gene was enriched in white adipocytes, induced early in differentiation and responded to dexamethasone, a typical glucocorticoid receptor (GR, encoded by *NR3C1*) agonist. Silencing of *NR3C1* in human multipotent adipose-derived stem cells (hMADS) or deletion of GR in a transgenic conditional mouse model results in diminished *HAND2* expression, establishing that adipocyte HAND2 is regulated by glucocorticoids via GR *in vitro* and *in vivo*. Using a combinatorial RNAseq approach we identified gene clusters regulated by the GR-HAND2 pathway. Interestingly, silencing of *HAND2* impaired adipocyte differentiation in hMADS and primary mouse adipocytes. However, a conditional adipocyte *Hand2* deletion mouse model using Cre under control of the *Adipoq* promoter did not mirror these effects on adipose tissue differentiation, indicating that Hand2 was required at stages prior to *Adipoq* expression. In summary, our study identifies HAND2 as a novel obesity-linked adipocyte transcription factor, highlighting new mechanisms of GR-dependent adipogenesis in human and mice.

## Introduction

Adipocytes are specialized fat cells that have a variety of functions including nutrient buffering, endocrine regulation, and thermogenesis. Obesity, the excess accumulation of white adipose tissue (WAT), is characterized by adipocyte dysfunction and metabolic imbalance. Approaches that enhance adipocyte health or increase the number of functional adipocytes have beneficial effects on systemic metabolism. Hence, a detailed molecular understanding of adipose tissue biology and pathology is of great therapeutic value. The adipose organ is composed of several depots containing mixed types of adipocytes, influenced by sex, age, diet as well as other genetic and environmental parameters. In principal, there are 2 types of adipocytes: white adipocytes, able to store and release lipids while providing nutrients and secreting adipokines as well as thermogenic adipocytes (brown and beige), which additionally dissipate chemical energy from nutrients as heat [1].

Adipogenesis, the formation of adipocytes, is a complex process regulated by the interplay of transcription factors, metabolites and hormonal cues. The commitment of mesenchymal stem cells to becoming preadipocytes and adipogenesis *per se* occurs in waves of transcriptional programs. During the early phase, progenitor cells lose their proliferative potential and start to differentiate. Genes involved in development e.g. bone morphogenic proteins, Wnt or Hh (hedgehog) proteins play a critical role during the commitment of progenitor cells to the adipocyte lineage [2]. Upon entering growth arrest, preadipocytes differentiate into adipocytes through the activation of transcription factor cascades responsible for the expression of genes important for terminal adipocyte function. The transcription factor complex of peroxisome proliferator-activated receptor-g (PPARg) and CCAAT/enhancer-binding protein-a (CEBPa) is a critical driver of adipocyte differentiation [3–5]. These and other transcription factors are triggered by a large panel of metabolites including dietary fatty acids or vitamins. Small molecules from the family of the thiazolidinediones (e.g. rosiglitazone) are PPARy ligands and strong promoters of adipocyte differentiation as well as adipocyte maturation [6]. Chronic rosiglitazone treatment has also been shown to induce a more thermogenic phenotype in preadipocytes isolated from WAT [7].

Hormones are broadly implicated in the metabolic derangements of adipose tissue function in obesity. Glucocorticoids (GCs) are steroid hormones modulating insulin sensitivity, blood glucose, lipid levels, and other metabolic parameters [8–10]. They are ligands for the glucocorticoid receptor (GR), a nuclear hormone receptor transcription factor. While GR regulates adipogenesis, lipolysis, and lipogenesis as well as thermogenesis [11], recent studies have demonstrated that GR is not required for the development of white or brown adipose tissue in mouse models [12–15]. However, the effect of GCs and GR on adipose tissue biology are multifaceted and depend on e.g. plasma concentrations, type of fat depot, sex, species, and the metabolic status of the individual. *In vivo*, physiological levels of GC mobilize energy under acute stress like the “fight or flight” response or store energy in the post-prandial phase when insulin levels increase [16]. In obesity, GCs play an important role in modulating inflammation, for example in adipose tissue by decreasing proinflammatory cytokine expression and macrophage infiltration [11]. Chronically elevated levels of GCs, either by pharmacologic treatment or in patients with Cushing’s syndrome results in partial lipodystrophy [17]. However, neither the adipose-specific activation of GR by GCs nor their regulation in obesity are fully understood.

In search of novel adipose-specific mechanisms linked to human obesity we have previously investigated methylation and gene expression signatures of visceral white adipose tissue (visWAT) and subcutaneous white adipose tissue (scWAT) and found that the transcription factor HAND2 (heart and neural derivatives expressed 2) to be among the top differentially expressed genes in human adipose tissue [18]. HAND2 encodes a helix-loop-helix transcription factor that has been described to play a role in stem cell differentiation into cardiomyocyte-like cells in cardiac morphogenesis [19]. This ability of HAND2 to control organ development and cell differentiation led us to hypothesize that HAND2 might be involved in adipocyte differentiation and function.

Here we show that HAND2 is an obesity-linked adipocyte transcription factor regulated by GCs and required for adipocyte differentiation *in vitro*. HAND2 was differentially expressed in adipose tissue depots with the highest levels in visWAT and inversely correlated to obesity in humans and mice. HAND2 expression was regulated by GC-GR signaling *in vivo* and *in vitro*. While HAND2 loss of function in human multipotent adipose-derived stem (hMADS) cells [20, 21] resulted in impaired adipocyte differentiation, mature adipocyte-specific knockout of HAND2 in mice was insufficient to affect energy homeostasis and other metabolic parameters. In summary, our study identifies HAND2 as a novel obesity-linked adipocyte transcription factor, highlighting new mechanisms of GR-dependent adipogenesis in human and mice.

## Results

### Adipose HAND2 is correlated to obesity in mice and men

We have recently identified *HAND2* amongst the top 3 differently methylated and expressed genes in visWAT versus scWAT of obese patients [18]. In order to investigate the importance of HAND2 in adipose tissue biology, we first determined its gene expression in different adipose depots obtained from group of obese and non-obese patients over a broad range of BMI. *HAND2* mRNA levels were higher in visWAT compared to scWAT (Figure 1 A) and scWAT depots showed substantially higher *HAND2* expression than human brown adipose tissue (BAT) (Figure 1 B). Intriguingly, *HAND2* expression in visWAT but not in scWAT was inversely correlated with BMI in these patients (Figure 1 C, D). However, *HAND2* in both tissues was correlated with body weight (Supplementary table 1). In line with these human data, adipose *Hand2* mRNA expression profiling of wild-type mice revealed that expression was markedly higher in gonadal WAT (gWAT), followed by scWAT and BAT. Moreover, gWAT *Hand2* was significantly lower in mouse models of obesity, including the high-fat diet (HFD)-induced obesity (DIO) as well as the genetically obese, leptin receptor-deficient *db/db* models (Figure 1 E, F), and also inversely correlated with body weight (Figure 1 G, H). These data demonstrate that Hand2 was prominently expressed in WAT depots and correlated with obesity in both mice and humans.

**Figure 1:**
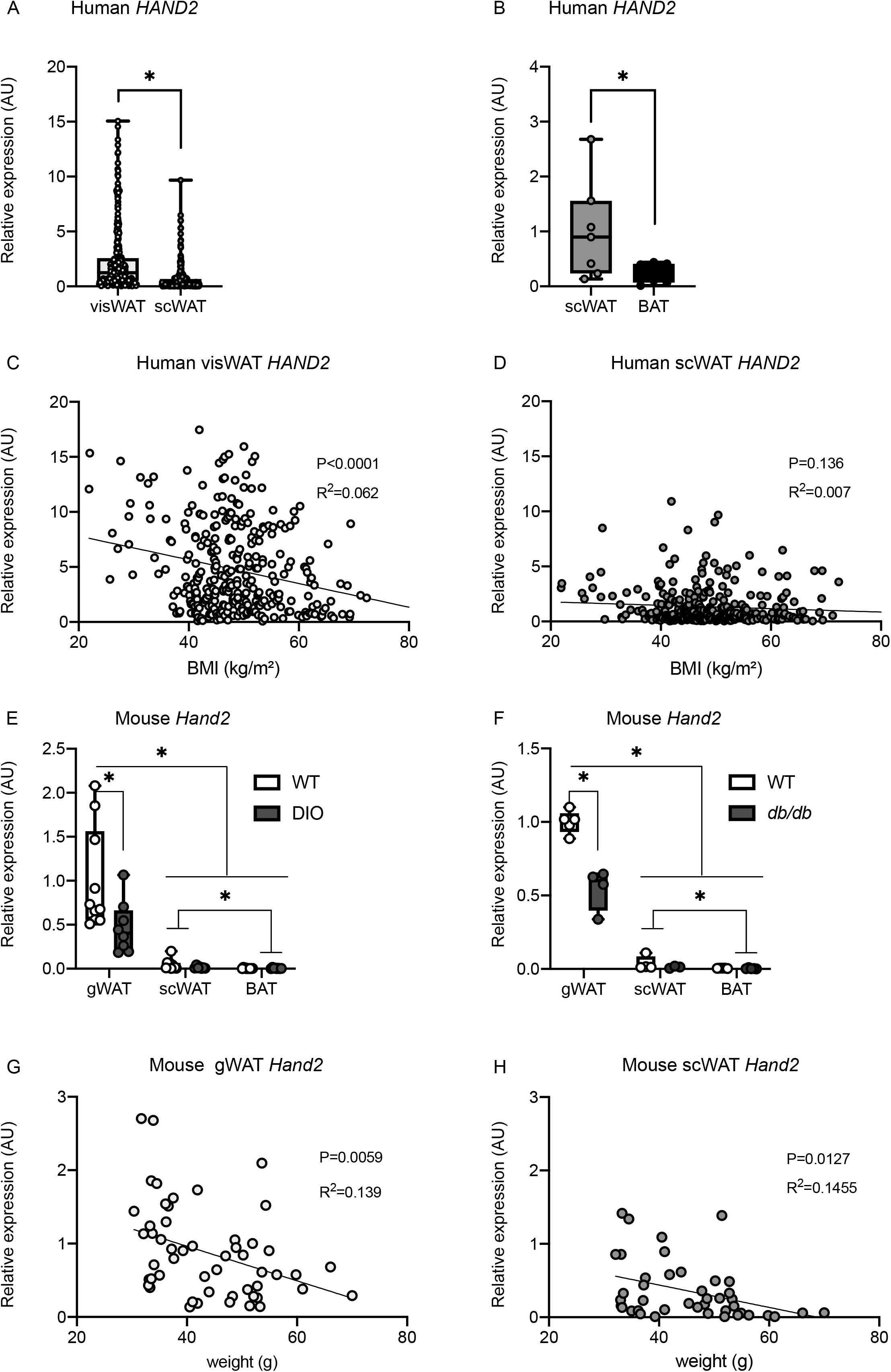
White adipocyte HAND2 is correlated to obesity in mice and men. *HAND2* expression in human visWAT vs scWAT (n=318) (A), and in human scWAT vs BAT (n=7) (B). *HAND2* expression correlated with BMI in visWAT (C) and scWAT (D) (n=318). *Hand2* expression in WT vs DIO mice (n=10) (E) and in WT vs *db/db* mice (n=5) (F). *Hand2* expression correlated with body weight in mouse gWAT (G) and scWAT (H) (n=54). Stat: two-tailed unpaired t-test (A), two-tailed paired t-test (B), correlation (C, D, G, H), two-way ANOVA (E, F). Statistical significance is indicated by *p<0.05.

### HAND2 is expressed in adipocytes and is induced early in adipogenesis

The cellular composition of adipose tissue is heterogenous and changes dynamically in obesity. To analyze the origin of Hand2 expression in more detail, we next measured *Hand2* mRNA levels in fractionated adipose tissue from mouse and human samples *ex vivo*. While mouse *Hand2* was poorly expressed in the macrophage fraction, its levels were higher in the stromal vascular fraction (SVF) compared to the adipocyte fraction (AF) (Figure 2 A, B). In line with these findings, *HAND2* levels tended to be higher in the SVF compared to the AF of human adipose tissue (Figure 2C). These findings indicate that Hand2 was predominantly expressed in preadipocytes and adipocytes. To further evaluate this concept, we employed *in vitro* adipocyte models. We first cultured primary adipocytes differentiated from SVF of different mouse adipose depots. *Hand2* expression was increased between D0 (day of induction of differentiation) and D3 (Figure 2 D). Differentiated adipocytes derived from the SVF of distinct anatomical locations mirrored the *in vivo Hand2* mRNA expression pattern with highest levels in adipocytes from gWAT compared to those isolated from scWAT or from BAT (Figure 2 E).

**Figure 2:**
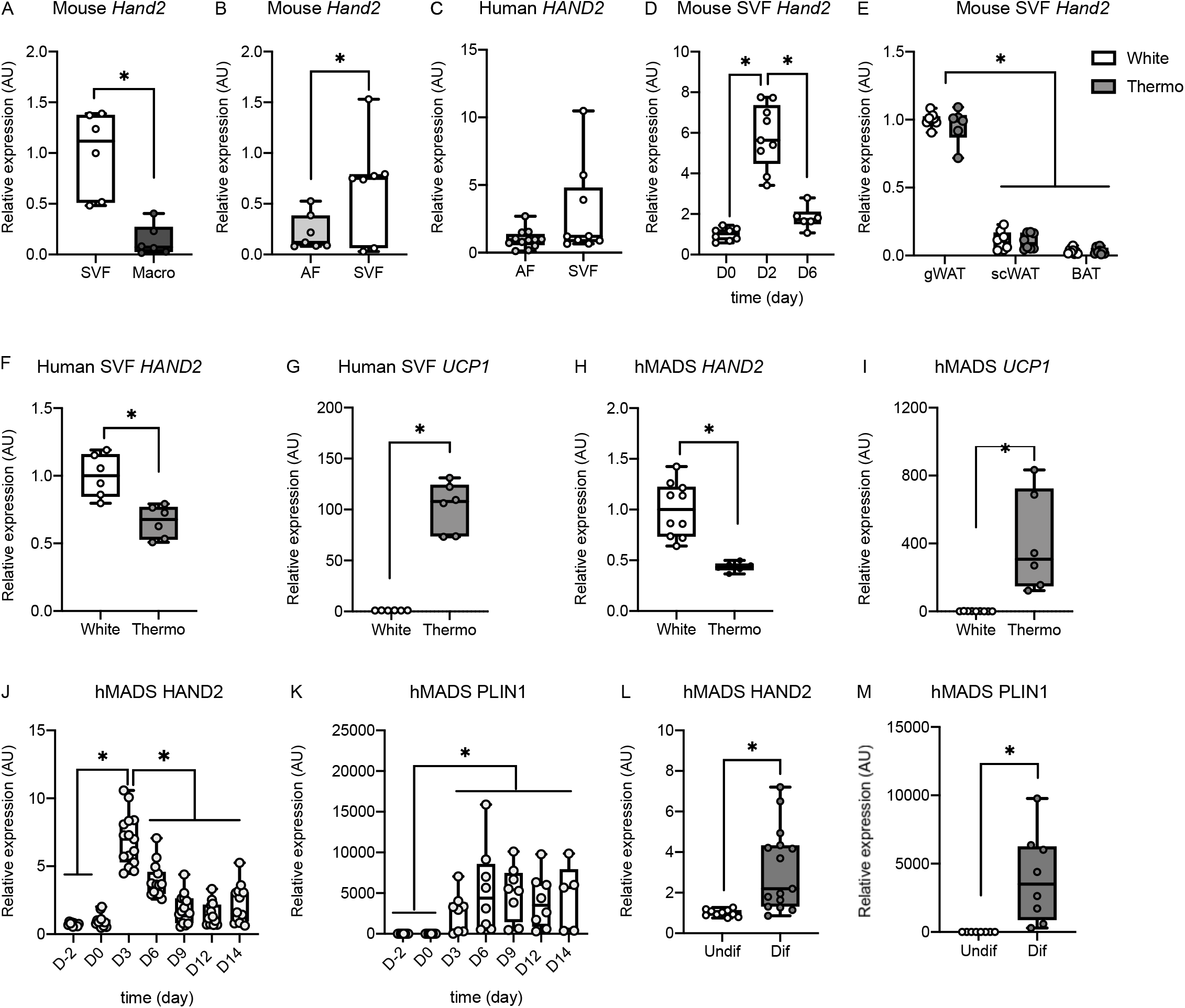
HAND2 is induced in early differentiation and expressed in both preadipocytes and mature adipocytes. *Hand2* expression in mSVF vs macrophages (A), vs Adipose Fraction (AF) (B) and in hAF vs SVF (C). *Hand2* expression during mSVF differentiation (D) and in mSVF from different fat depos differentiated in white or thermogenic adipocytes (E). *HAND2* (F) and *UCP1* (G) expression in hSVF differentiated in adipocytes. *HAND2* (H) and *UCP1* (I) expression in hMADS mature adipocytes. *HAND2* (J) and *PLIN1* (K) expression in hMADS cells during differentiation. *HAND2* (L) and *PLIN1* (M) expression in hMADS preadipocytes and mature adipocytes. Stat: two-tailed unpaired t-test (A, B, C, F, G, H, I, L), one-way ANOVA (D, J, K), two-way ANOVA (E). Statistical significance is indicated by *p<0.05.

Interestingly, chronic rosiglitazone treatment, which induces thermogenic differentiation [7], did not affect *Hand2* expression in mouse cells (Figure 2 E). In adipocytes differentiated from human adipose SVF, *HAND2* was higher in the classical white compared to the rosiglitazone-induced thermogenic differentiation regimen (Figure 2 F, G). The hMADS cell model has been described as a reliable tool for studying the metabolism of white and thermogenic adipocytes [20, 21]. Also, in hMADS cells, *HAND2* was higher in the white compared to the thermogenic differentiation regimen (Figure 2 H, I). Furthermore, we confirmed that mRNA *HAND2* was induced early in hMADS adipogenesis (Figure 2 J, K). Interestingly, fully differentiated hMADS adipocytes showed a slightly higher expression of HAND2 than preadipocytes (Figure 2 L, M). In summary, these data illustrate that HAND2 is highly and selectively expressed in white adipocytes with a spike of expression during the commitment phase towards the adipocyte lineage.

### Loss of HAND2 impairs adipogenesis

As HAND2 was induced in early adipogenesis in human and mouse adipocyte models, we hypothesized that HAND2 might be an important component of the adipogenesis program. To test this hypothesis, we silenced *HAND2* gene expression in hMADS cells before the induction of differentiation (Figure 3 A). *HAND2*-specific siRNA treatment led to diminished levels of *HAND2* mRNA at D0 and D2 compared to control siRNA-treated cells (Figure 3 B). Transcriptomic analysis at D2 revealed that silencing of *HAND2* was associated with a marked downregulation of the expression of prominent mature adipocyte genes, including *ADIPOQ, APOE, LIPE* and *PLIN1* (Figure 3 C). Also, key adipogenic transcription factors such as *PPARG, CEBPA* and *PPARGC1B* were expressed at lower levels (Figure 3 D-F), indicating that HAND2 was required for proper execution of the adipogenic program. Ingenuity pathway analysis confirmed that silencing of *HAND2* led to a broad dysregulation of the transcriptional programs required for adipocyte biology, e.g. insulin receptor signaling, fatty acid β-oxidation, and triacylglycerol biosynthesis (Supplementary figure 1 A, B). Interestingly, among the *in silico* predicted inhibited upstream regulators were *NR3C1* (encoding the glucocorticoid receptor, GR), *KLF15* and *PPARG*, all well-known master regulators of adipogenesis. In contrast, upstream regulators associated with proliferation such as *FGF2, TGFB* or *MIF* were predicted to be activated (Supplementary figure 1 C). The aberrant execution of the transcriptional adipogenesis program was also mirrored in the overall cellular phenotype, as early *HAND2* silencing completely abolished the differentiation of hMADS cells into mature adipocytes as demonstrated by the lack of lipid droplet formation (Figure 3 G, H). Also, in mouse SVF-derived preadipocytes, early silencing of *Hand2* led to lower lipid droplet formation while the induction of key adipogenic genes, including *Plin1* and *Pparg* remained largely intact (Figure 3 I, J, K, L, M). Taken together, our results indicate that Hand2 was required for adequate differentiation of both mouse and human white adipocytes.

**Figure 3:**
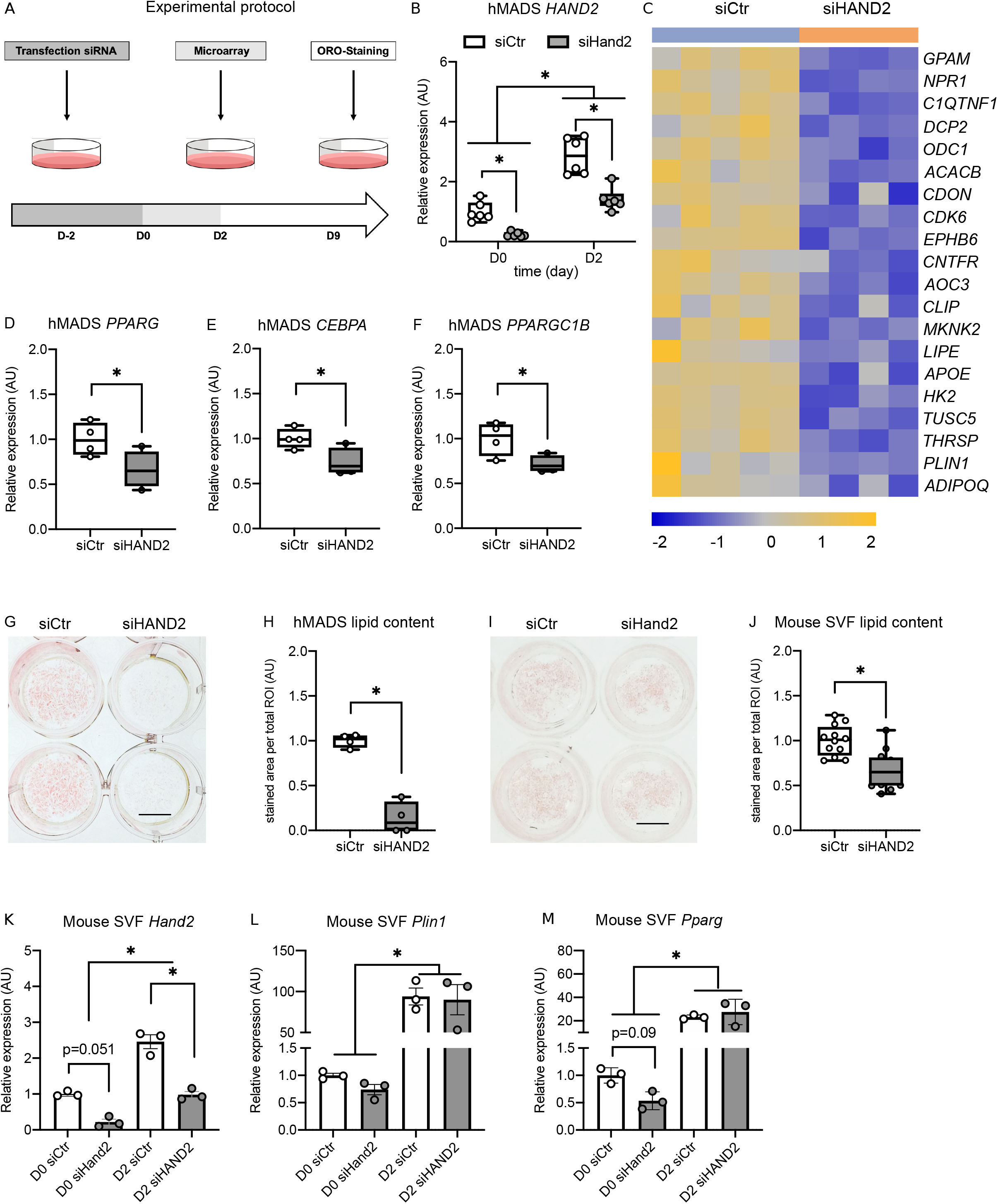
Loss of HAND2 impairs adipocyte differentiation. hMADS cells were transfected with siHAND2 before induction of the differentiation. RNAs were collected at day 0 and day 2 or cells were differentiated until day 9 and stained with Oil Red O. Day 2 samples were analyzed by microarray (A). *HAND2* expression at day 0 and day 2 (B). Heat map, generated from the microarray data, of the most inhibited genes (C). Gene expression of adipogenesis markers (D, E, F). Oil Red O staining of hMADS cells (G, H) and mSVF (I, J) differentiated. *Hand2, Plin1* and *Pparg* expression in mSVF transfected with siHand2 at day −2 and collected at day 0 and day 2. Stat: two-way ANOVA (B), two-tailed unpaired t-test (D, E, F), one-way ANOVA, mean +/- SEM (K, L, M). unpaired t-test (H, J). Statistical significance is indicated by *p<0.05.

### Hand2 in mature adipocytes is dispensable in vivo

In order to assess whether Hand2 might also be required for adipocyte function and systemic metabolic control *in vivo*, we created a conditional Cre-loxP mouse model for adipocyte-specific deletion of Hand2. We used an established transgenic mouse model, in which critical parts of the *Hand2* gene are flanked by loxP sites [22] and crossed this model with mice carrying Cre driven by the *Adipoq* promoter [23], which led to genetic deletion of *Hand2* in mature adipocytes *in vivo* (*Hand2*^AdipoqCre^) (Supplementary figure 2 A, B). As adipocyte function is an important pillar of lipid homeostasis during fasting and the postprandial phase, we first tested plasma triglycerides and non-esterified fatty acid levels in chow-fed male and female mice. Fasted and refed plasma lipid levels remained largely unchanged between *Hand2*^AdipoqCre^ and wild-type littermate controls (Supplementary figure 2 C-F). These observations were supported with human metabolic data showing no correlation between *HAND2* expression in WAT and plasma triglyceride levels. (Supplementary table 1) Also, adipocytes are critically important for sustaining metabolic control in states of caloric excess, so we next tested the role of adipocyte *Hand2* in DIO. Both male and female mice were placed on HFD for 12 weeks to induce weight gain and insulin resistance whereas chow diet-fed mice were included as controls (Figure 4 A). As expected, HFD feeding induced markedly higher weight gain compared to chow diet in both sexes, whereas the absence of Hand2 in white adipocytes did not have an effect (Figure 4 B, C). During the study, we performed glucose tolerance tests after 6 and 12 weeks as well as an insulin tolerance test after 12 weeks of the feeding regimen. As expected, HFD-fed mice displayed markedly lower glucose tolerance as well as markedly higher insulin resistance relative to chow-fed controls at both time points, however, we did not detect any genotype-specific differences (Figure 4 D-I). At the end of the study, while lean mass remained unchanged across diets and genotypes, HFD-fed mice displayed markedly higher adiposity and WAT depots weights than chow-fed controls and these differences were independent of sex and genotype (Figure 3 J, S). A careful analysis of adipose tissue histology revealed the expected increase in adipocyte diameter in WAT upon HFD feeding, but overall WAT and BAT were phenotypically similar when comparing genotypes (Supplementary figure 3 A-F). In summary, while *Hand2* was required for a proper adipogenesis *in vitro*, genetic deletion of *Hand2* in mature adipocytes *in vivo* using *Adipoq*-Cre did not impact DIO and the development of insulin resistance.

**Figure 4:**
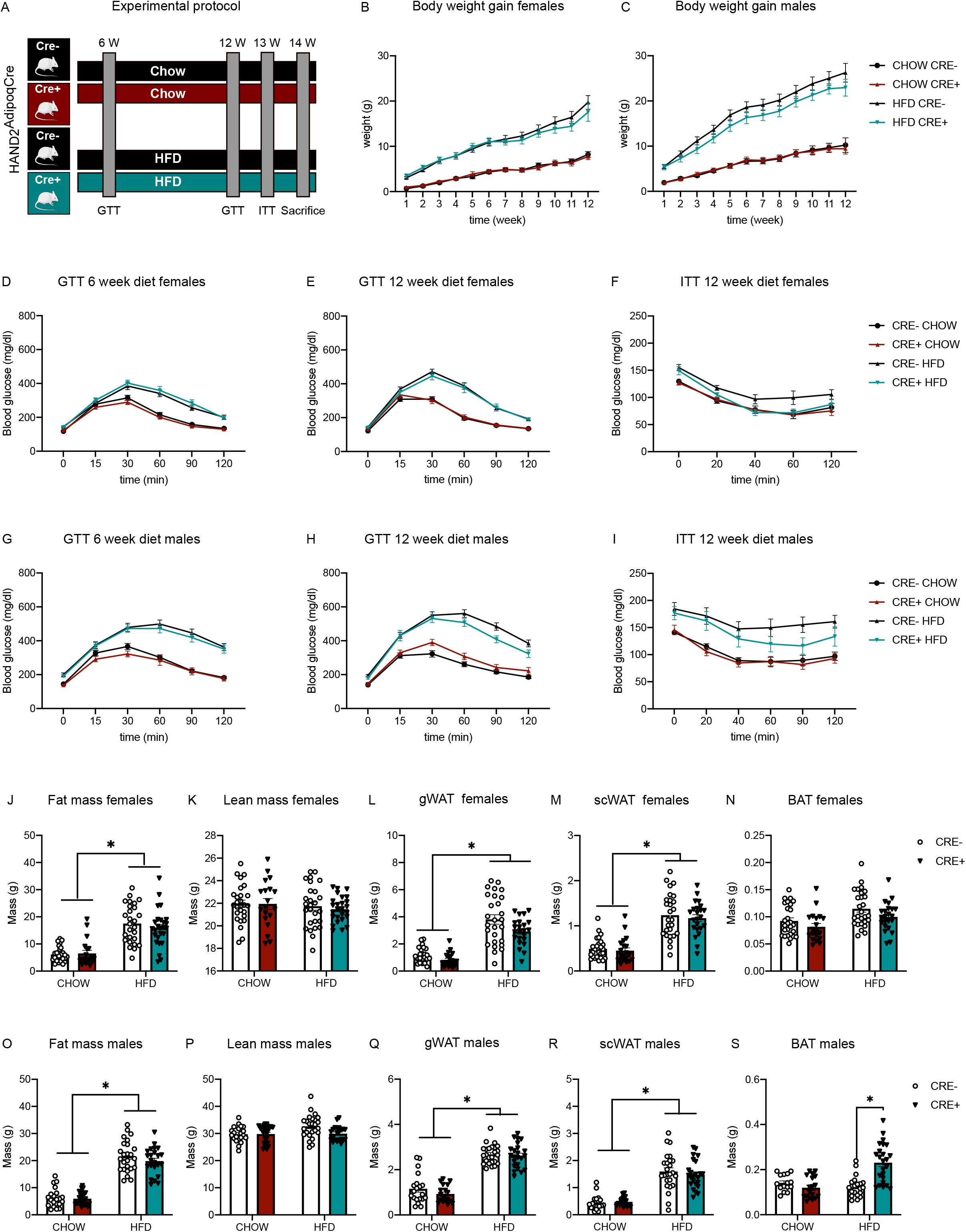
Metabolic phenotyping of HAND2^AdipoqCre^ mice fed an HFD. *Hand2*^AdvpoqCre^ (CRE+) and WT (CRE-) littermates, females and males, were fed an HFD (60%) for 6 or 12 weeks. Several metabolic parameters were measured (A). Body weight gain in females (n = 20 to 28) (B) and males (n = 21 to 26) (D). GTT in females (D) and males (G) after 6 weeks of HFD diet. GTT and ITT in females (E, F) and males (H, I) after 12 weeks of HFD diet. Final measurement of body composition (J, K, O, P) and adipose depos weight (L, M, N, Q, R, S) in males and females; Stat: two-way ANOVA; mean +/- SEM (C-E). Statistical significance is indicated by *p<0.05.

### HAND2 is regulated by glucocorticoids via the glucocorticoid receptor

Our results supported the hypothesis that Hand2 expression levels in adipocytes were determined early during adipogenesis, and that Hand2 expression was required for differentiation of stem cells into adipocytes *in vitro*. Among the most powerful and established inducers of adipogenesis are insulin, cAMP-raising agents (e.g. isobutylmethylxanthine, IBMX), agonists of PPARg (e.g. rosiglitazone), and GR agonists, such as dexamethasone (DEX). As *HAND2* expression was induced in cell culture upon treatment with this commonly used adipogenic hormone cocktail, we explored the ability of the individual ingredients to regulate *HAND2* expression. We found that DEX, but not the other adipogenic agents, induced *HAND2* expression (Figure 5 A). This induction of *HAND2* by DEX was antagonized by simultaneous treatment with the GR antagonist RU-486 in hMADS cells (Figure 5 B, C) as well as in SVF cells (Figure 5 D, E) irrespectively of whether it was added before or after differentiation. Of note, as expected perilipin gene expression was very low in preadipocytes, markedly higher in differentiated adipocytes, and was unaffected by pharmacological manipulation of GR. Also *in vivo*, a single DEX injection led to higher *Hand2* expression in gWAT (Supplementary figure 4 A, B). Chronic DEX treatment induced an increase of energy expenditure in the light phase and a decrease in the dark phase, nevertheless no specific phenotype was observed in *Hand2*^AdipoqCre^ mice (Supplementary figure 4 C-E). Other metabolic parameters including activity, food intake, body weight, body composition and blood glucose concentration were not influenced by the absence of Hand2 in mature adipocytes, neither at 22 °C nor at 30 °C thermoneutrality (Supplementary figure 4 F-M). Congruent with a GR-dependent mechanism, siRNA-mediated knockdown of *NR3C1* in hMADS preadipocytes as well as mature adipocytes (Figure 6 A, B) and genetic inactivation in SVF from gWAT of Hand2^flox/flox^ mice with Cre-encoding mRNA both in preadipocytes as well as mature adipocytes (Figure 6 C, D) abolished the effect of DEX on *HAND2*. Using a similar approach with preadipocytes isolated from WAT of *Nr3c1*^flox/flox^ mice, allowing the genetic deletion of GR [24], we found that genetic deletion of GR abrogated the increase in *Hand2* (Figure 6 E). For inhibition of *Nr3c1* in differentiated adipocytes we used several models. First, we silenced *Nr3c1* in differentiated adipocytes from SVF by RNAi and observed that loss of *Nr3c1* diminished the increase in *Hand2* expression (Figure 6 F). Of note, in all the conditions described above, as expected, *PLIN1* expression was almost inexistent in preadipocytes compared to adipocytes and remained unaffected by the different treatments (Supplementary figure 5). Also, we used *Nr3c1*^flox/flox^ mice crossed with mice expressing the global tamoxifen-inducible Cre-ERT2 fusion protein (GR^ERT2CRE^) [25]. In SVF cells from these animals, we confirmed that GR regulated *Hand2* expression as tamoxifen treatment abolished the DEX response (Figure 6 G), altogether supporting the notion that HAND2 was regulated by the GC/GR pathway during adipocyte differentiation.

**Figure 5:**
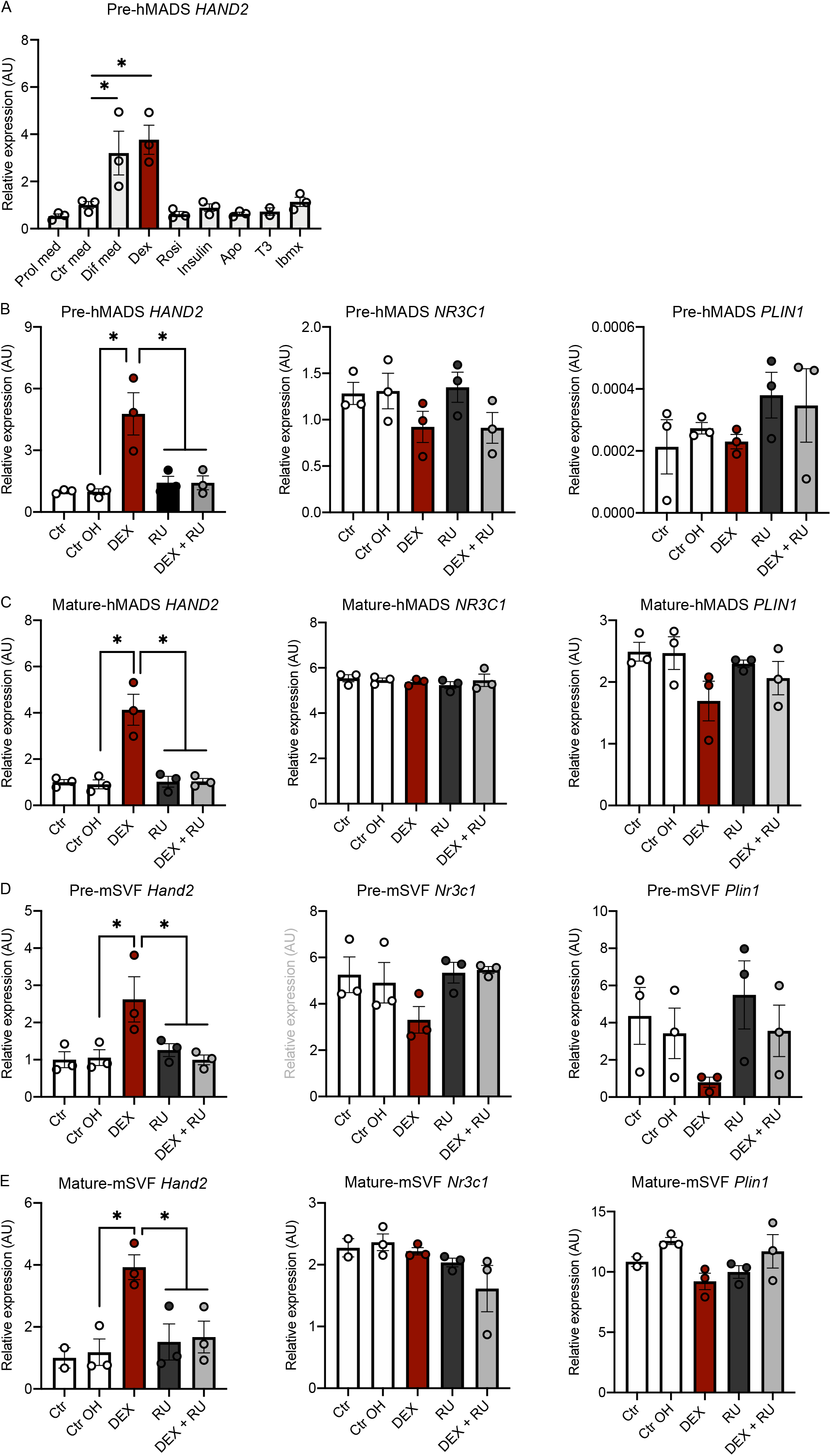
Dexamethasone regulates HAND2 expression. Testing for all single components of our standard differentiation cocktail in terms of *HAND2* mRNA induction. Prol med (proliferation media), Ctr media (differentiation media D0 without any components), Dif med (differentiation media D0 with all components) DEX, Rosi, Insulin, Apo, T3, IBMX (differentiation media at D0 with only one component) (A). *HAND2, NR3C1* and *PLIN1* expression in hMADS preadipocytes (B) and mature adipocytes (C) treated with DEX or/and RU-486. *Hand2, Nr3c1* and *Plin1* expression in mSVF preadipocytes (D) or mature adipocytes (E) treated with DEX and/or RU-486. Stat: two-tailed unpaired t-test; mean +/- SEM. Statistical significance is indicated by *p<0.05.

**Figure 6:**
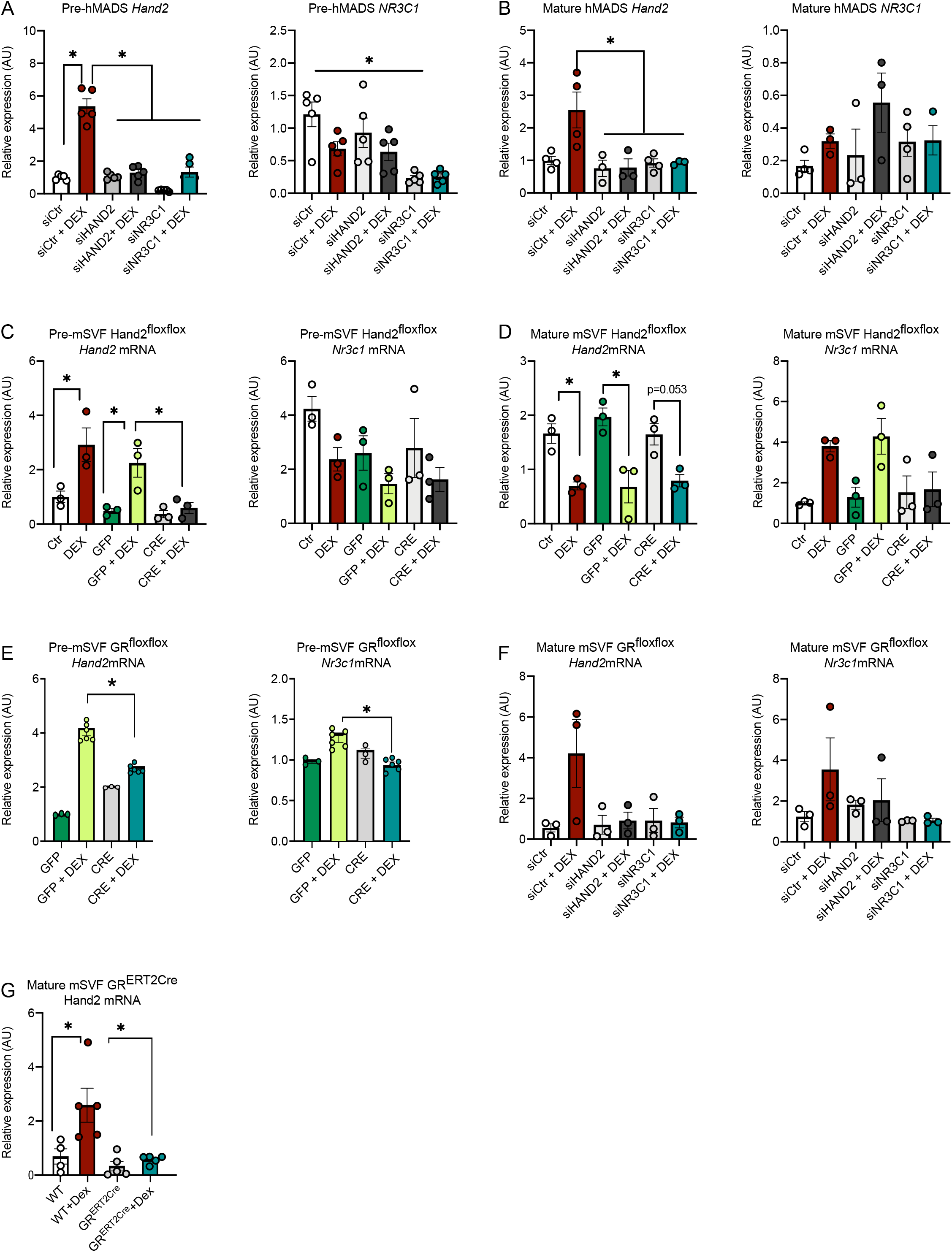
HAND2 is regulated by glucocorticoids via the glucocorticoid receptor. *HAND2* and *NR3C1* expression in hMADS preadipocytes (A) and mature adipocytes (B) transfected with siHAND2 or siNR3C1, treated or not with DEX. *Hand2* and *Nr3c1* expression in mSVF preadipocytes (C) and mature adipocytes (D) from HAND2^flox/flox^ mice, as well as mSVF preadipocytes from GR^flox/flox^ (E) transfected either with a mRNA GFP or CRE recombinase treated or not with DEX. *Hand2* and *Nr3c1* expression in mSVF differentiated in adipocytes and transfected with siHAND2 or siNR3C1 treated or not with DEX. *Hand2* expression in adipocytes differentiated from mSVF from GR^ERT2CRE^ mouse treated with tamoxifen and treated or not with DEX (G). Stat: one-way ANOVA; mean +/- SEM (A-G). Statistical significance is indicated by *p<0.05.

### Gene networks during adipocyte differentiation regulated by the GR-HAND2 pathway

Our results indicated that HAND2 was induced by GR and while HAND2 was required for adipocyte differentiation it was dispensable in mature adipocytes. Therefore, we set out to understand the transcriptional networks during early adipogenesis downstream of the GR-HAND2 pathway. Using RNAi, we silenced *HAND2* and *NR3C1* in hMADS cells, either alone or in combination, treated the cells 2 days later with DEX for 12 hours and performed RNAseq (Figure 7 A, B, C). A principal component analysis revealed that the transcriptome signature of cells with *NR3C1* silencing and treated with DEX clustered together with the control samples without DEX treatment (Figure7 D), illustrating the global requirement of GR for the DEX effects on adipogenesis. However, cells with *HAND2* silencing clustered together with control cells treated with DEX, suggesting HAND2 regulates only a small subset of GR-related genes. Using a gene set enrichment analysis [26, 27], we determined the transcriptional networks regulated by GC/GR signaling for benchmarking our data set. Indeed, we found many confirmed GC/GR signaling pathways, underlining the reliability of our dataset (Figure 7 E). Next, we focused on gene expression levels that were differentially and commonly regulated by GR and HAND2 as part of the GR-HAND2 pathway. In this unbiased and global analysis, we found four main pathways to be regulated. The metapathway biotransformation phase I and II, including genes from the cytochrome p450 family implicated in the synthesis of cholesterol steroids and other lipids. Interestingly, the mTORC1 pathway and the large family of class A rhodopsin-like GPCRs were also found. Lastly, the growth factor signaling pathway VEGFA-VEGFR2 was identified as a potential downstream effector pathway (Figure 7 F). In summary, these data confirm that HAND2 plays an important role very early in the adipocyte differentiation process and might play an important role in the execution of the GC/GR program.

**Figure 7:**
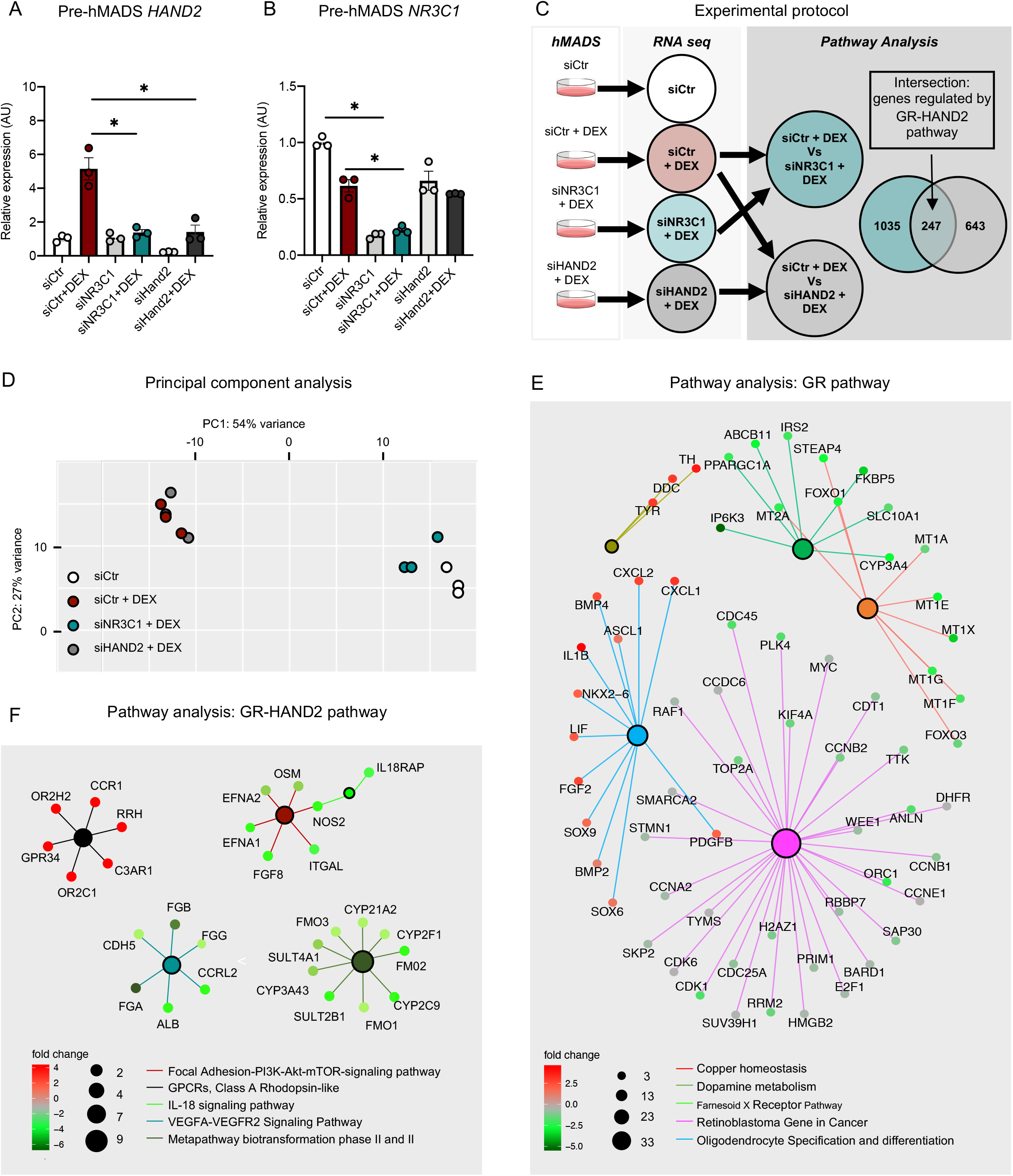
Gene networks during adipocyte differentiation regulated by GR-HAND2pathway. *HAND2* (A) *NR3C1* (B) expression in hMADS cells 48h after transfection siNR3C1 or siHAND2 treated or not with DEX. RNAs samples were analyzed by RNAseq (C). Principal component analyses of the RNAseq data (D). GO pathway analysis of siCtr + DEX vs siNR3C1 + DEX (E) and of the intersection between siCtr + DEX vs siNR3C1 + DEX and siCtr + DEX vs siHAND2 + DEX (F). Stat: one-way ANOVA; mean +/- SEM (A, B). Statistical significance is *p< 0.05.

## Discussion

The ability of adipocytes to safely store extra calories in the form of lipids is a critical component of a healthy metabolism. Understanding the transcriptional mechanisms regulating adipocyte differentiation and adipose tissue expansion is fundamental to our understanding of basic metabolic principles as well as for developing novel obesity therapeutics. In this study, we show that HAND2 is a critical factor for early adipogenesis, regulated by GC/GR signaling and correlated to body weight and obesity in mice and men. In human and mouse cell culture models, loss of *HAND2* in stem cells or preadipocytes impaired adipocyte differentiation. The earlier we manipulated *HAND2* expression, the more severe was the effect in interfering with proper adipocyte differentiation: in hMADS cells it completely abolished differentiation while in preadipocytes isolated from SVF, which are already committed to becoming adipocytes, the effects were similar but much smaller in magnitude. Therefore, it should not come as a surprise that our conditional *Hand2* mouse model using *Adipoq*-Cre did not show any overt metabolic phenotype, as the *Adipoq* gene and hence the Cre recombinase, are expressed relatively late in adipocyte differentiation. Indeed, adiponectin gene expression appears after the commitment phase and is almost absent from SVF cells [23]. One way to probe this further would be to deplete *Hand2* under a Cre driver specially expressed in the commitment phase of the adipocytes, such as Wnt or Hedgehog [28, 29]. Nevertheless, those master regulators are not specific to adipocytes and might lead to lethality in very early stage of the embryo development similarly to the global deletion of *Hand2* [30, 31]. A whole-body Cre approach using the aforementioned tamoxifen-inducible model [32] has similar legitimate limitations.

The second major finding of our study is that *HAND2* expression is regulated by GCs via GR during early and late stages of adipogenesis. DEX is an established adipogenic agent used in most *in vitro* adipogenesis models. We found that DEX induced *HAND2* expression in the first days of differentiation. Our RNAseq results demonstrated that GR is required for the global effects of DEX, which was expected. In contrast, cells with loss of HAND2 still had a relatively intact transcription profile, indicating that HAND2 only regulates a small and defined set of genes upon DEX treatment. Further analyses are required to interrogate the relevance of these putative downstream effectors of HAND2 for adipogenesis. In addition, although DEX treatment has been described to boost adipocyte differentiation, GR is not essential for adipogenesis in mouse models [15]. However, the transcription factors *DEXRAS1* and *KLF15* have been described as direct targets of GR, affecting adipocyte differentiation [33, 34]. In the case of *KLF15*, in addition of being activated by GR its expression also depends on the expression of *CEBPβ* and *CEBPδ* [34]. In our microarray study, we identified *KLF15* as one of the potential adipogenic upstream regulators inhibited by loss of *HAND2*. Interestingly, it seems that there is crosstalk between mTORC1 and GR signaling via KLF15 in the context of glucocorticoid-induced muscle atrophy [35]. Even though HAND2 does not dramatically affect global gene expression regulated by DEX, we show that mTORC1-dependent genes are regulated by the GR-HAND2 pathway. Considering the pleiotropic effects of DEX and GR signaling in the body, it is possible that some of the previous functions attributed to HAND2, for example in the heart, also involve GR activity as well as vice versa that established GR-mediated effects for example in immune cells could also involve HAND2.

Lastly, we found that *HAND2* expression is inversely correlated with BMI both in human obesity as well as in mouse models of dietary and genetic obesity. Our study demonstrates that adipose *HAND2* is predominantly expressed in the adipocyte/preadipocyte moiety of the tissue, and only low levels are found in tissue-resident macrophages. This implies that the correlation of the *HAND2* expression in obesity reflects a down regulation of the adipocyte expression. It is possible that lowering of *HAND2* levels with increasing BMI limits adipogenesis. Reactivation of the *HAND2* gene could therefore help stimulating adipogenesis and healthy adipose tissue expansion, thus restoring insulin sensitivity and metabolic health. In summary, our study introduces HAND2 as a novel player in adipogenesis and highlights a new layer of GC/GR signaling, thus enhancing our understanding of adipocyte biology in obesity.

## Methods

### Additional methods can be found in the supplementary methods

#### Cell culture and stromal vascular fraction preparation

Human multipotent adipose-derived stem cells characterization has been previously described [20, 21]. Cells were grown and differentiated as previously described [36]. Stroma vascular fraction of mouse adipose depots from scWAT, gWAT and BAT was differentiated as previously described [37]. Human SVF was isolated from abdominal subcutaneous human adipose tissue, collected from healthy patients (abdominoplasty) and differentiated following the same protocol as hMADS cells. The study was approved by the University Ulm ethical committee (vote no. 300/16) and all patients gave written informed consent.

#### Gene expression and functional analysis *in vitro*

Preadipocytes and mature adipocytes (human and mouse models) were transfected with 20 nM of siCtr, siHAND2 or siNR3C1 (ON-TARGETplus siRNA smart pool,) or with 1 μg/ml of mRNA CRE or mRNA GFP (StemMACS Cre Recombinase mRNA). Chemical activation and inhibition of GR were performed by over-night treatment with respectively 100 nM Dexamethasone (D4902 Sigma-Aldrich) and/or 200 nM RU486 (M8046 Sigma-Aldrich). RU486 was added 3 hours before DEX. Lipid amount was determined by Oil Red O staining (Sigma O1391). Images were analyzed using the commercially available software Definiens Developer XD 2 (Definiens AG, Germany). We used the % Oil Red O stained area per total region of interest (ROI). ROI was defined by the surface of the well. RNA were extracted using TRIzol reagent as previously described [36]. All the experiments have been done at least 3 times. qPCR analyses were performed using SYBR Green master mix. The sequence of the probes used are reported in (Supplementary table 2).

#### Human studies

Study protocol applying for gene analysis of human BAT versus scWAT was reviewed and approved by the ethics committee of the Hospital District of Southwestern Finland, and subjects provided written, informed consent following the committee’s instructions. The study was conducted according to the principles of the Declaration of Helsinki. All potential subjects were screened for metabolic status, and only those with normal glucose tolerance and normal cardiovascular status (as assessed on the basis of electrocardiograms and measured blood pressure) were included. The age range of the subjects was 23–49 years. We studied a group of 7 healthy volunteers (2 men and 5 women). BAT was sampled from positive FDG-PET scan sites in supraclavicular localization, and subcutaneous WAT was derived via the same incision. The human scWAT versus visWAT samples were collected in the context of a cross-sectional study of 318 individuals (249 women, 69 men; BMI range: 21.9 – 97.3 kg/m^2^, age range: 19-75 years) Abdominal omental and subcutaneous WAT samples were collected during elective laparoscopic abdominal surgery as described previously [38]. Adipose tissue was immediately frozen in liquid nitrogen and stored at −80 °C. The study was approved by the Ethics Committee of the University of Leipzig (approval no: 159-12-21052012) and performed in accordance to the declaration of Helsinki. All subjects gave written informed consent before taking part in this study. Measurement of body composition and metabolic parameters was performed as described previously [38, 39].

#### Mouse experiments

All animal studies were conducted in accordance with German animal welfare legislation and protocols were approved by the state ethics committee and government of Upper Bavaria (nos. ROB-55.2-2532.Vet_02-16-117 and ROB-55.2-2532.Vet_02-17-125). All mice were maintained in a climate-controlled environment with specific pathogen-free conditions under 12-h dark–light cycles in the animal facility of the Helmholtz Center Munich. *Hand2* expression in WT versus DIO mice was performed in C57BLK/6J males 18 weeks old (10 animals per group). *Hand2* gene expression correlated with body weight included 54 wild type C57BLK/6J and NMRI male mice (18 to 24 weeks old). Mice were fed ad libitum with regular rodent chow. Constitutive adipose tissue specific Hand2 knockout mice (*Hand2^AdipoqCre^*) were generated by crossing *Adipoq*^CRE^ mice (Jackson laboratory, stock number 028020; C57BLK/6J) with *Hand2*^floxflox^ mice kindly given by the laboratory of professor Rolf Zeller (NMRI) [22]. *Hand2^AdipoqCre^* (CRE+) and wild type littermates (CRE-) were used for all experiments. Animals were fed an HFD 60% (D12492, Research Diets Inc., New Brunswick, NJ, USA) ad libitum from the age of 6 weeks for 12 weeks. Glucose and insulin tolerance tests were performed after 6 and 12 weeks of high-fat feeding, respectively. Animals were fasted at 8 am for 4 to 6 hours and subsequently injected intraperitoneally with glucose at 2g/kg or insulin 0.8 U/kg (HUMINSULIN Normal 100, Lilly Germany, Bad Homburg). Blood samples were taken from the tail vein before and 15, 60, 90 and 120 min after the i.p. injection to assess glucose levels. (AccuChek Performa glucose meter, Roche Diabetes Care, Mannheim, Germany Assessment of the fat mass vs lean mass was performed using whole-body magnetic resonance analysis (EchoMRI, Houston, TX). The experiment has been performed on 3 different cohorts of littermates with a total of 20 to 28 animals per group.

Adult male and female NMRI (wild type, 10 week of age) received an acute intraperitoneal injection of either DEX (1 mg/kg, USP grade; Millipore Sigma) or vehicle (2% ethanol) at 8 am. Animals were exposed to DEX treatment for 6h. At termination, the animals were killed by cervical dislocation and organs were collected and immediately frozen in liquid nitrogen and stored at −80 °C. The experiment has been done twice with a total of 7 to 10 animals per group.

For indirect calorimetry, *Hand2^AdipoqCre^* (CRE+) and wild type female littermates (CRE-) were acclimatized to the TSE PhenoMaster cages (TSE Systems, Bad Homburg, Germany) for 1 week (experiment performed once with 6 animals per group). Basal indirect calorimetry analysis, including energy expenditure, food consumption, oxygen consumption and locomotor activity, was performed for 3 days. For three subsequent days the animals were injected with DEX in IP (1mg/kg) every morning at 8 am.

## Acknowledgement

We thank Ez-Zoubir Amri (Institut de Biologie Valrose, Université Nice Sophia Antipolis, France) for providing hMADS cells, as well as Rolf Zeller (Developmental Genetics, Department of Biomedicine, University of Basel, 4058 Basel, Switzerland) for providing the *Hand2*^floxflox^ mice. We thank Anne Loft, Anastasia Georgiadi and Pauline Morigny for discussion and Mareike Bamberger for excellent technical assistance. M.G. was supported by Alexander von Humboldt Foundation postdoctoral fellowships. P.F.P. was supported by the Deutsche Forschungsgemeinschaft (FI 1700/7-1, Heisenberg program). J.T. was supported by the Deutsche Forschungsgemeinschaft (TU 220/13-1). A.B. was supported by the Deutsche Forschungsgemeinschaft Sonderforschungsbereich 1123 (B10), and the Deutsches Zentrum für Herz-Kreislauf-Forschung Junior Research Group Grant. S.H. and J.B. were supported by the Helmholtz Future topic ‘Aging and Metabolic Programming, AMPro’. S.H. was supported by the Deutsche Forschungsgemeinschaft Trans-Regio TRR205 and the Deutsches Zentrum für Herz-Kreislauf-Forschung Standortprojekt Cardiometabolism.

The authors declare no conflicts of interest.

## Authors contribution

Conceptualization: M.G., M.S., A.B. and S.H.

Methodology: M.G.

Formal Analysis: M.G., F.F.T.

Investigation: M.G., F.F.T., G.C, S.K, E.S.V., M.I, C.G., M.G.L., S.K., D.H, M.R.G, A.M, A.F, M.B.

Mouse material donation: J.T., A.A.D. and F.M.

Human material donation: D.T., P.F.P., M.W., M.B. and K.V.

Writing: M.G., A.B. and S.H.

Supervision: S.H., A.B., M.S., J.B.

Project administration: M.G.

Funding Acquisition: M.G., S.H., A.B., J.T., J.B.

