## Supplementary material for "HAND2 is a novel obesity-linked adipogenic transcription factor regulated by glucocorticoid signaling"

Supplementary table 1: Correlation analysis between *HAND2* expression in visWAT and scWAT and clinical data.

| | *HAND2* | | | |
|---|---|---|---|---|
| | visWAT | | scWAT | |
| | R | P | R | P |
| ***Anthropometric data*** | | | | |
| Body weight (kg) | -0.1963 | **0.0004** | -0.1164 | **0.0386** |
| BMI (kg/m²) | -0.2545 | **<0.0001** | -0.083 | 0.1368 |
| ***Metabolic data*** | | | | |
| FPG (mmol/l) | -0.1137 | **0.0433** | -0.004 | 0.9429 |
| FPI (pmol/l) | -0.032 | 0.5682 | -0.1732 | **0.002** |
| HOMA-IR | -0.06214 | 0.2716 | -0.1622 | **0.0039** |
| Total cholesterol (mmol/l) | -0.03918 | 0.6683 | 0.2444 | **0.0064** |
| HDL-C (mmol/l) | -0.03354 | 0.714 | 0.03375 | 0.7121 |
| LDL-C (mmol/l) | -0.03896 | 0.674 | 0.1869 | **0.041** |
| TG (mmol/l) | -0.09665 | 0.2876 | 0.01051 | 0.9078 |

**Supplementary table 2: Sequence primer for** SYBR Green **qPCR.**

| Gene | sequence |
|---|---|
| m_Hand2_F | TCCAAGATCAAGACACTGCG |
| m_Hand2_R | TCTTCTTGATCTCCGCCTTG |
| m_Nr3c1_F | AGCTCCCCCTGGTAGAGAC |
| m_Nr3c1_R | GGTGAAGACGCAGAAACCTTG |
| m_Plin1-F | CAAGCACCTCTGACAAGGTTC |
| m_Plin1-R | GTTGGCGGCATATTCTGCTG |
| m_Tbp_F | ACCCTTCACCAATGACTCCTATG |
| m_Tbp_R | ATGATGACTGCAGCAAATCGC |
| m_Hprt_F | TCAGTCAACGGGGGACACATAAA |
| m_Hprt_R | GGGGCTGTACTGCTTAACCAG |
| h_HAND2_F | GACACTCCCGTGTGGTAAGG |
| h_HAND2_R | AAGGGGTTGAGTAGGTTGGC |
| h_PLIN1_F | ACCCCCCTGAAAAGATTGCTT |
| h_PLIN1_R | GATGGGAACGCTGATGCTGTT |
| h_NR3C1_F | AAAGAGACGAATGAGAGTCCTTGGA |
| h_NR3C1_R | GCTTGCAGTCCTCATTCGAGTTT |
| h_TBP_F | ACGCCAGCTTCGGAGAGTTC |
| h_TBP_R | CAAACCGCTTGGGATTATATTCG |
| h_36B4_F | TGCATCAGTACCCCATTCTATCAT |
| h_36B4_R | AGGCAGATGGATCAGCCAAGA |

Supplementary figure: 1

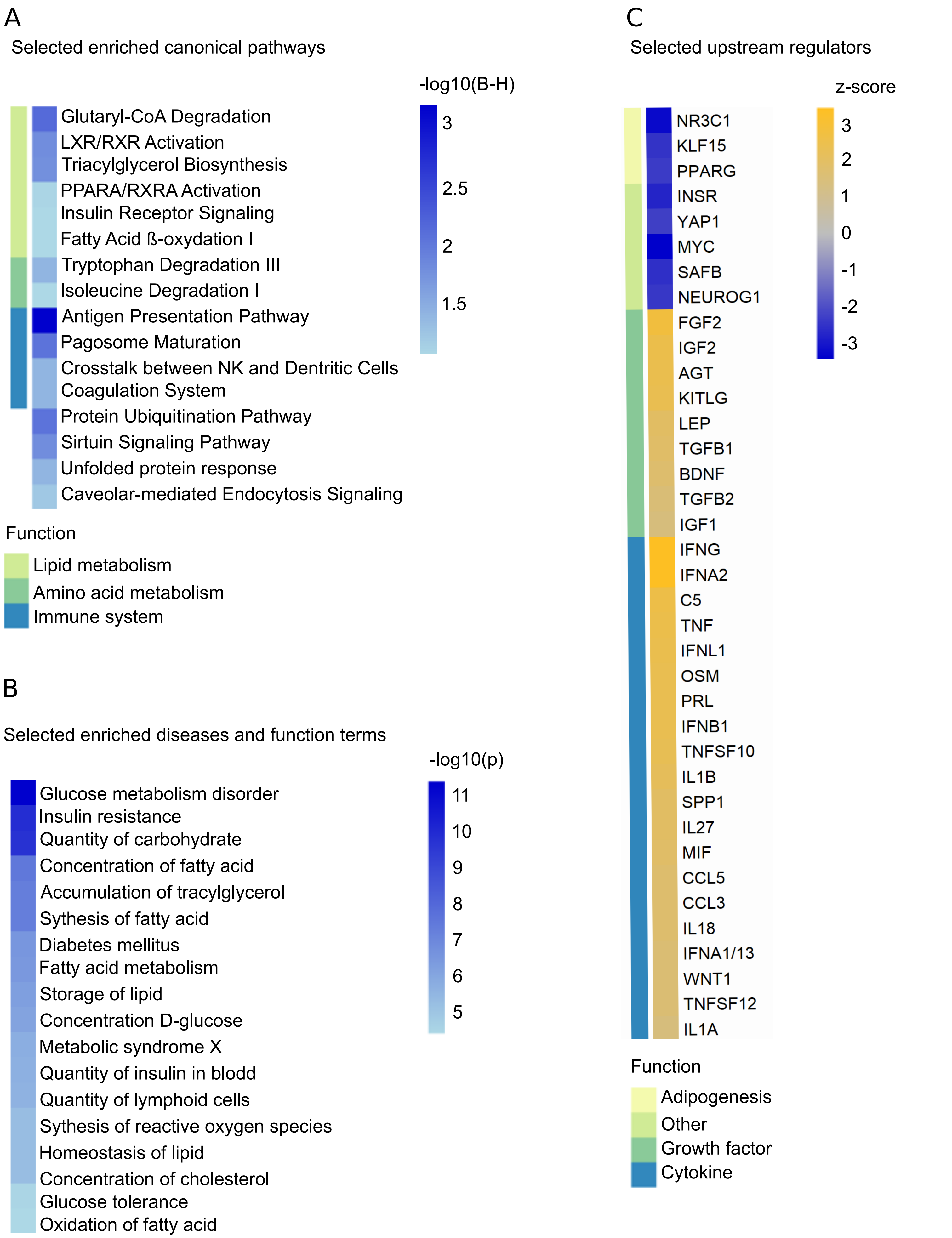

### Supplementary figure: 2

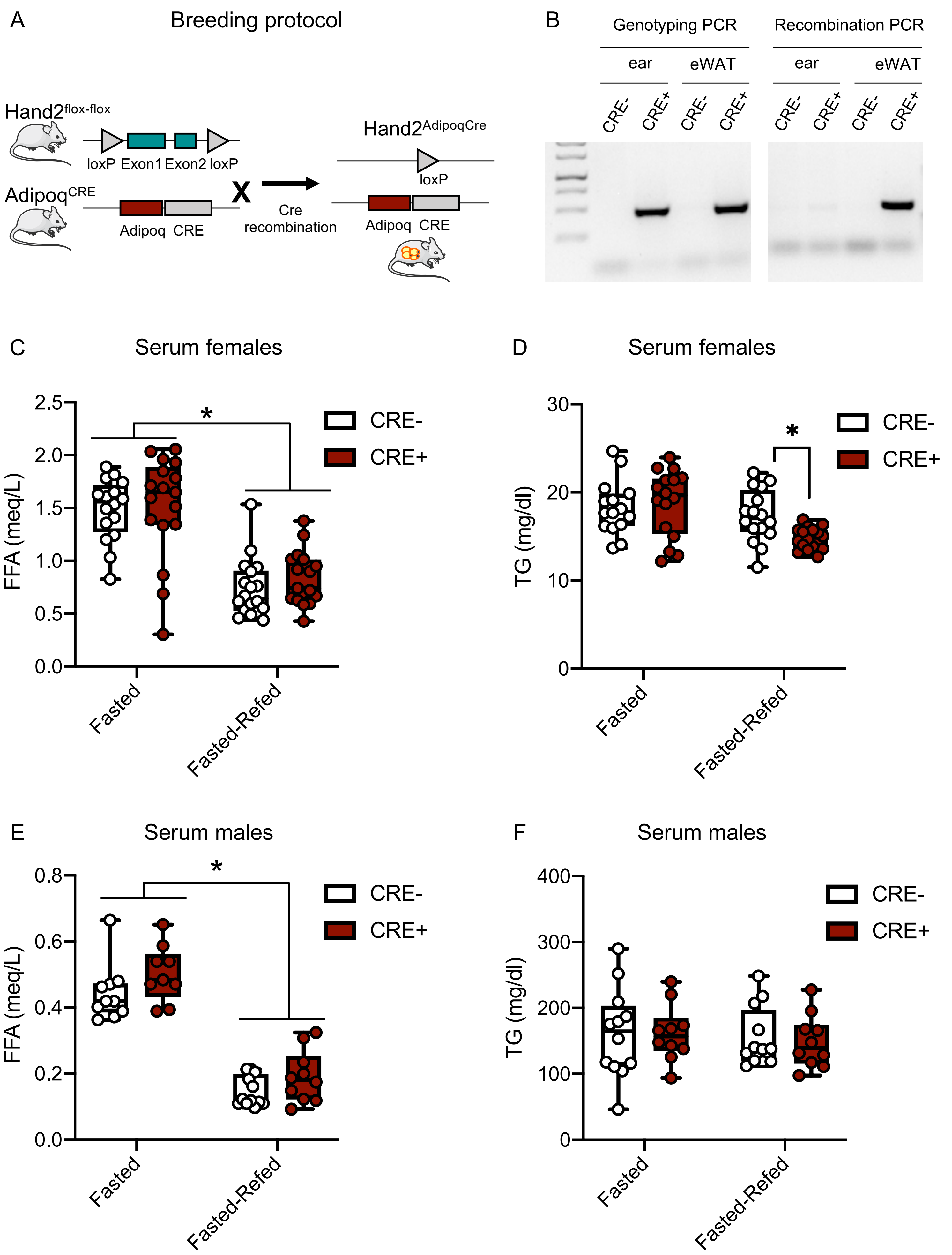

### Supplementary figure: 3

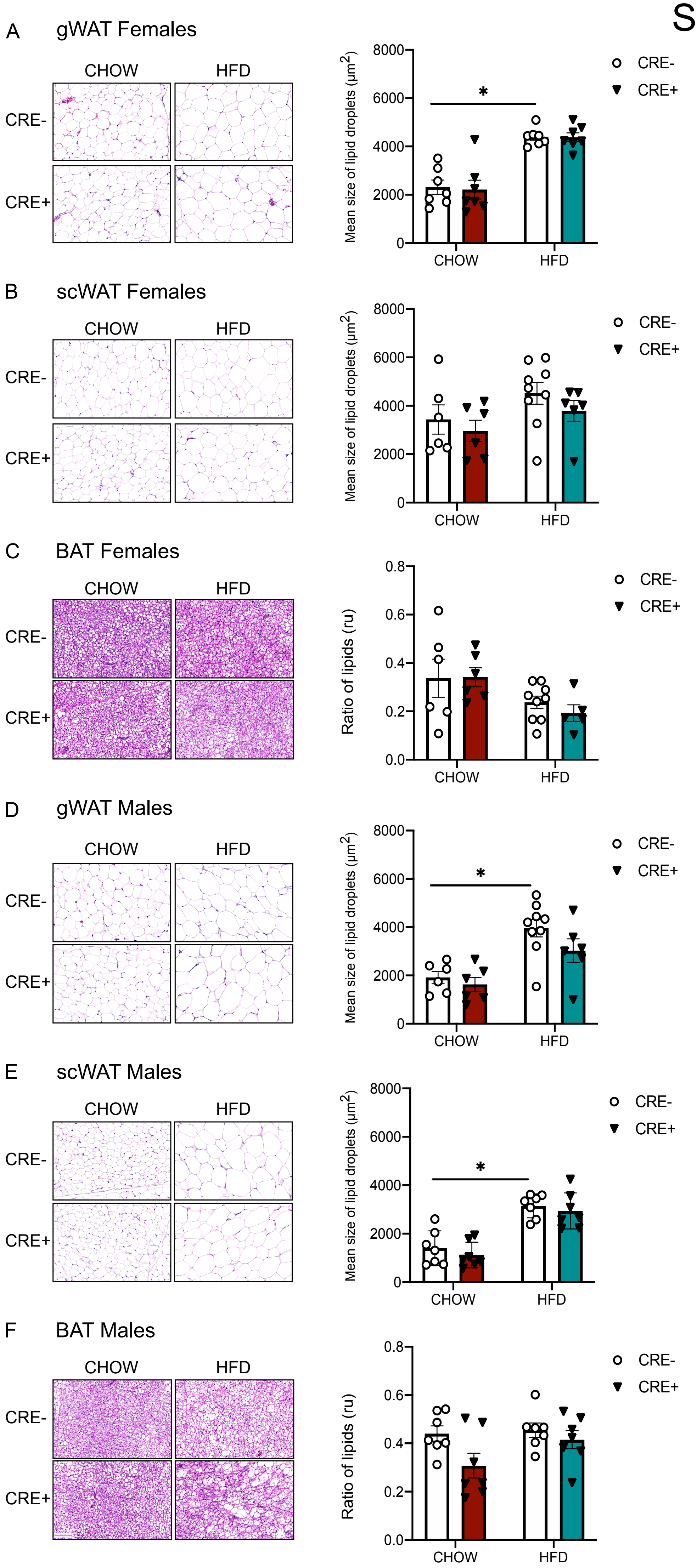

### Supplementary figure: 4

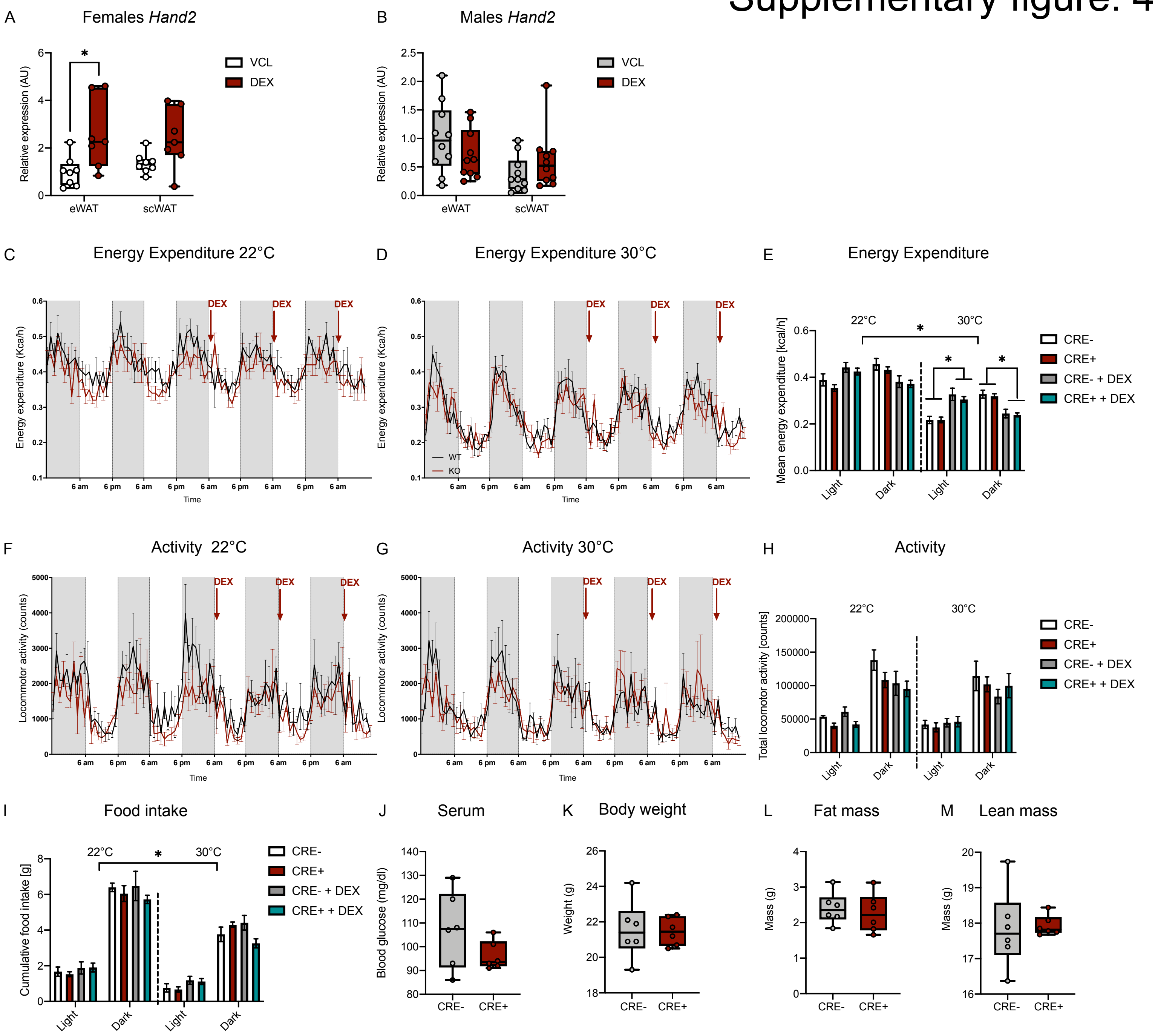

### Supplementary figure: 5

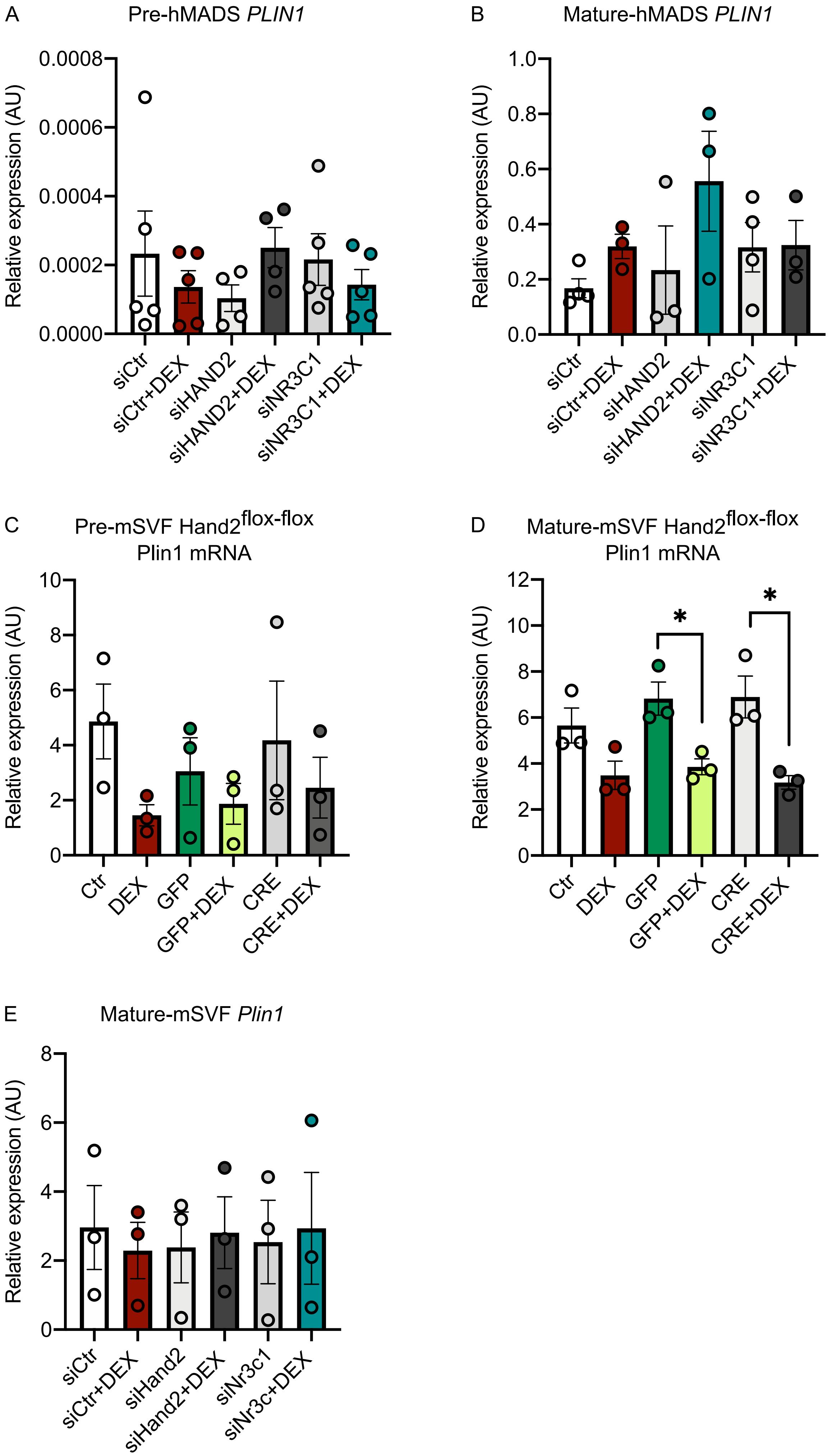

***Supplementary legends:***

***Supplementary table 1: Correlation analysis between HAND2 expression in visWAT and scWAT and clinical data.***

HAND2 expression in human visWAT vs scWAT (n=318) correlated with anthropometric and metabolic data.

Stat: Pearson correlation coefficient was computed. R (Pearson r), P (p value).

***Supplementary figure 1: Ingenuity Pathway Analysis of microarray data of hMADS cells loss of function for HAND2***

Significantly regulated genes (FDR<10%) from the analysis siCtr vs siHAND2 were used as input and enriched terms from the Canonical Pathways analysis (A) and Disease&Functions analysis (B) are shown. Predicted significantly activated (z-score >2) or inhibited (z-score <-2) upstream regulators are shown in (C). Terms and regulators were selected mainly for relevance to metabolism. All growth factors and cytokines in the Upstream Regulator analysis had z-scores >2 and are shown.

***Supplementary figure 2: Metabolic phenotyping of $HAND2^{AdipoqCre}$ mice.***

Breeding protocol of the *$HAND2^{AdipoqCre}$* mouse line (A). *Hand2* DNA expression illustrating the ubiquitous expression of CRE recombinase and the adipose tissue specific recombination of *Hand2* (B). FFA and TG levels in the serum of *$HAND2^{AdipoqCre}$* and wild type littermate females (n=17) and males (n=10 to 12) after 12 hours fasting or 6 hours fasting followed by 12 hours refeeding.

Stat: one-way ANOVA, (C-E). Statistical significance is indicated by *p<0.05.

***Supplementary figure 3: Histological analysis of $HAND2^{AdipoqCre}$ mice fed an HFD.***

An eosin/hematoxylin staining was performed on histological cut of the different adipose tissues of WT or *$HAND2^{AdipoqCre}$* mice fed a CHOW or an HFD for 12 weeks. Pictures were taken and the size of the adipocytes was measured in females gWAT (A), scWAT (B), BAT (C), and in males gWAT (D), scWAT (E), BAT (F).

Stat: two-way ANOVA; mean +/- SEM (C-E). Statistical significance is indicated by *p<0.05.

***Supplementary figure 4: Metabolic phenotyping of** $HAND2^{AdipoqCre}$ **mice acutely injected with dexamethasone***

Wild type animals were injected with DEX and scarified 6 hours later. *Hand2* expression was measured in gWAT and scWAT of females (A) and males (B). Females *HAND2*$^{AdipoqCre}$were housed in metabolic cages. Energy expenditure (C-E), activity (F-H) and food intake (I) were measured over a period of 5 days at 22°C or thermoneutrality, 30°C, including 3 daily injection with DEX. Serum (J), body weight (K), fat mass (L) and lean mass (M) were recorded before the experiment.
Stat: two-way ANOVA; mean +/- SEM (A, B, E, H, I); two-tailed unpaired t-test (J-M). Statistical significance is indicated by *p<0.05.

***Supplementary figure 5: Expression of PLIN1 in hMADS cells and hSVF presenting a loss of function for HAND2 or NR3C1.***

*PLIN1* expression in hMADS preadipocytes (A) and mature adipocytes (B) transfected with siHAND2 or siNR3C1; in mSVF preadipocytes (C) and mature adipocytes (D) from HAND2$^{flox/flox}$ mice; in mSVF differentiated in adipocytes and transfected with siHAND2 or siNR3C1.
Stat: one-way ANOVA, mean +/- SEM (A-E). Statistical significance is indicated by *p<0.05.

**Supplementary methods:**

*Transcriptomic analyses:*

Microarray expression profiling: Total RNA was isolated employing the RNAeasy kit (Qiagen) using the small and large RNA protocol for animal tissue and cultured cells, including on-column DNase digestion. The Agilent 2100 Bioanalyzer was used to assess RNA quality and only high-quality RNA (RIN>7) was used for microarray analysis. Total RNA (150 ng) was amplified using the WT PLUS Reagent Kit (Thermo Fisher Scientific Inc., Waltham, USA). Amplified cDNA was hybridized on Human Clariom S arrays (Thermo Fisher Scientific). Staining and scanning (Gene Chip Scanner 3000 7G) was done according to manufacturer`s instructions. Samples were processed in two batches; 2$^{nd}$ batch was: Control_8/10, siRNA_Hand2_9/12). Transcriptome Analysis Console (TAC; version 4.0.0.25; Thermo Fisher Scientific) was used for quality control and to obtain annotated normalized SST-RMA gene-level. Statistical analyses were performed by utilizing the statistical programming environment R (Rainer et al. 2006; R Development Core Team 2010) .

Genewise testing for differential expression was done employing the limma t-test including batch correction and Benjamini-Hochberg (BH) multiple testing correction (FDR < 10%). To reduce background, gene sets were filtered for TAC Data Above Background p-values<0.05. Heat maps were generated in R. Pathway analyses were generated through the use of Ingenuity Pathway Analysis software (IPA®, QIAGEN Redwood City, www.qiagen.com/ingenuity) using Fisher`s Exact Test p-values (Diseases&Functions), BH-corrected p-values (Canonical Pathways), or z-scores (Upstream Regulators).

Array data has been submitted to the GEO database at NCBI (GSE148699).

RNA-seq library construction and sequencing: Library preparation and sequencing were performed by Novogene. An amount of 800ng of total RNA was used and controlled quantitatively and qualitatively with Agilent Bioanalyzer 2100 system. Sequencing libraries were generated using NEBNext® Ultra™ RNA Library Prep Kit for Illumina® (NEB, USA) following manufacturer’s recommendations. Prepared PCR products were purified and size-selected on AMPure XP beads (Beckman A63881). The prepared libraries were sequenced (paired-end) on an Illumina platform._Read counts post processing, statistical analysis and differential gene expression analysis were performed using DESeq2 (Anders and Huber 2010) in R (R Development Core Team 2010). In order to observe the overall effect of experimental covariates a two-dimensional PCA plot was employed. GO gene enrichment analysis of differentially expressed genes was performed using ClusterProfiler (Yu et al. 2012).

*Analysis of HAND2 mRNA expression in human adipose tissue:*

Human scWAT vs viscWAT: RNA from adipose tissue was extracted by using RNeasy Lipid tissue Mini Kit (Qiagen, Hilden, Germany). Quantity and integrity of RNA was monitored with NanoVue plus Spectrophotometer (GE Healthcare, Freiburg, Germany). 1µg total RNA from SC and Vis adipose tissue and liver were reverse-transcribed with standard reagents (Life technologies, Darmstadt, Germany). cDNA was then processed for TaqMan probe-based quantitative real-time polymerase chain reaction (qPCR) using the QuantStudio 6 Flex Real-Time PCR System (Life technologies, Darmstadt, Germany). Expression of HAND2 was calculated by standard curve method and normalized to the expression of hypoxanthine guanine phosphoribosyltransferase 1 (HPRT1) as a housekeeping gene. The probes (Life technologies, Darmstadt, Germany) for HAND2 (Hs01047149_m1 and Hs00232769_m1) and HPRT1 (Hs01003267_m1) span exon-exon boundaries to improve the specificity of the qPCR.

*Histology:*

The samples were fixed in 4% (w/v) neutrally buffered formalin and subsequently routinely embedded in paraffin. 3 µm thick sections were stained with hematoxylin and eosin (HE), using a HistoCore SPECTRA ST automated slide stainer (Leica, Germany) with prefabricated staining reagents (Histocore Spectra H&E Stain System S1, Leica, Germany), according to the manufacturer's instructions. Stained tissue sections were scanned with an AxioScan. Z1 digital slide scanner (Zeiss, Jena, Germany) equipped with a 20x magnification objective. Quantification of lipid amount was morphometrically determined by automatic digital image analysis using the commercially available software Definiens Developer XD 2 (Definiens AG, Germany). The calculated parameter for BAT was the ratio of total area of lipid droplets per whole tissue section and for WAT the mean size of lipid droplets per whole tissue section.
